# Identifying High-Dimensional Genomic Factors Modulating Biological Networks Across Multi-Omics Data

**DOI:** 10.64898/2025.12.18.695273

**Authors:** Samuel Anyaso-Samuel, Shilan Li, Giovanny Herrera Ossa, Emily Vogtmann, Xiaoyu Wang, Xing Hua, Fei Qin, Wei Zhao, Mohammad L. Rahman, Xiaohong R. Yang, Kevin Brown, Bin Zhu, Steven C. Moore, Christian Abnet, Tongwu Zhang, Maria Terresa Landi, Kai Yu, Paul S. Albert, Jianxin Shi

**Author notes:** **Correspondence to:** Paul S. Albert, Ph.D., Senior Investigator, Biostatistics Branch, Division of Cancer Epidemiology and Genetics, National Cancer Institute, Jianxin Shi, Ph.D., Senior Investigator, Biostatistics Branch, Division of Cancer Epidemiology and Genetics, National Cancer Institute.

## Abstract

Biological traits such as genes, metabolites, and microbial taxa interact within complex networks, yet how genomic factors shape these interactions remains poorly understood. Here, we introduce GFBioNet, a computationally efficient method for identifying factors that modulate direct associations between biological traits within network models. Our two-stage strategy first estimates a baseline network using Gaussian graphical models and then tests whether genomic factors modulate specific network edges (trait–trait relationships), enabling scalable analysis of high-dimensional multi-omics data while explicitly controlling the false discovery rate (FDR). Simulations demonstrate reliable FDR control and high statistical power across a broad range of settings. Applied to multiple datasets, GFBioNet reveals host genetic variants influencing oral microbiome relationships, gut microbial taxa modulating metabolite networks in colorectal cancer, and somatic mutations and copy-number alterations reshaping gene expression networks in lung adenocarcinoma. By expanding network analysis to evaluate modifiers of trait–trait relationships, GFBioNet offers a versatile tool for uncovering the genomic architecture of biological networks across multi-omics studies.

## Introduction

Gaussian graphical models (GGMs)^1-3^ are widely used to explore associations among numerous biological traits and construct networks across diverse application domains, such as gene expression^4^, microbial taxa^5-7^, metabolites^8^, and proteome^9^. Conceptually, GGMs aim to identify pairs of traits with non-zero partial correlation coefficients (PCCs), which represent direct dependencies between traits while accounting for the effects of all other traits. A biological network is thus constructed by connecting any two variables with PCC significantly different from zero.

Various statistical methods have been developed for building GGMs, which are generally grouped into three categories: estimating sparse precision matrices^2,3,10^, penalized nodewise regression^11^, and limited-order partial correlation methods^12-14^. These techniques have been refined to enhance sensitivity for specific applications, such as modeling shared or heterogeneous structure across conditions^15,16^, explicitly identifying hubs^17,18^, or combining both strategies^7,19^. The application of GGMs has yielded valuable insights, improving our understanding of biological traits and the etiology of diseases.

In this manuscript, we address a statistical problem of identifying high-dimensional factors that modify direct trait–trait dependencies within biological networks. Specifically, given a set of traits (e.g., gene expression levels in blood or tissue samples) and a set of modifying factors (e.g., germline common variants), our goal is to identify triplets (*i, j, t*), in which factor *t* modifies the PCC between traits *i* and *j*. For instance, in **Figure 1a** and **1b**, the PCC between the expression levels of *BTN2A1* and *BTN3A2* is associated with the genotype of a single nucleotide polymorphism (SNP) *rs1935234* in *CD*4_*NC*_ cells, based on single-cell transcriptome sequencing data from the OneK1K cohort^20^. This type of second-order association, examining how a factor influences the relationship between two traits rather than the traits individually, has been seldom explored but may reveal regulatory or mechanistic patterns not evident from marginal analyses.

**Figure 1.**
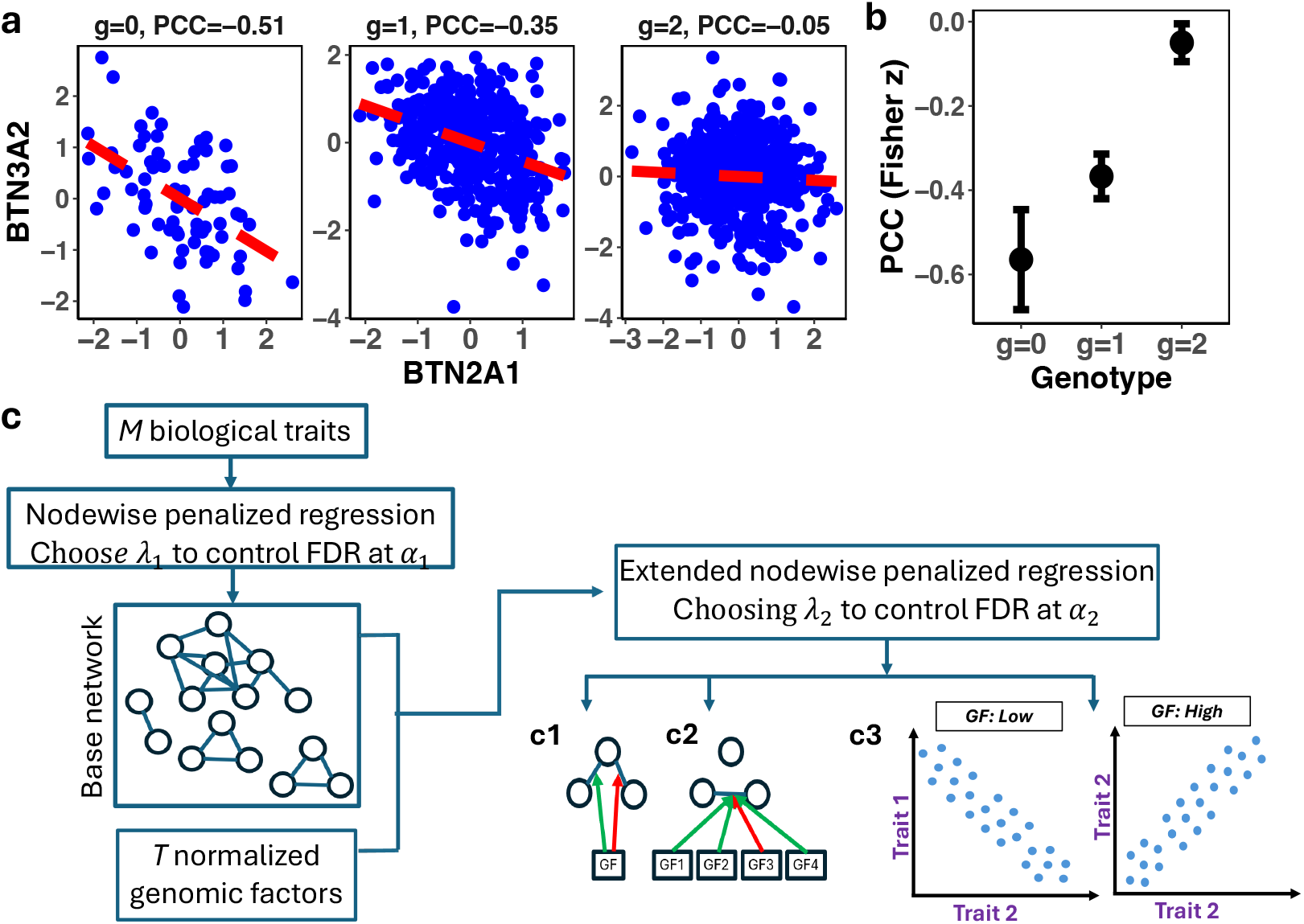
Overview of GFBioNet for identifying genomic factors (GF) associated with biological networks. (a) Partial correlation coefficients (PCCs) between expression levels of *BTN2A1* and *BTN3A2* increase with the genotypic value of *rs1935234* in CD4_NC_ cells, based on OneK1K single-cell transcriptome data. Scatter plots show gene expression values adjusted for other relevant genes. (b) Fisher’s transformation z values and their standard error in three genotype groups (copies of minor allele). (c) **GFBioNet, a** two-stage framework for identifying GFs influencing trait–trait networks. Stage 1: Construct a baseline network via node-wise penalized regressions; edges represent direct trait associations. Stage 2: Identify GFs that modify baseline edges by modeling GF–trait interactions, with false discovery rate (FDR) control applied at each stage. (c1) One GF modulates multiple trait–trait interactions (green = positive, red = negative modifications). (c2) Multiple GFs modulate a single network edge. (c3) Example scatter plots stratified by GF values (low/high for continuous GFs), illustrating identified signals.

We introduce a statistical framework, GFBioNet (**G**enomic **F**actors modulating **Bio**logical **Net**works), designed to identify factors that influence direct dependencies between traits within a GGM framework using penalized nodewise regression. In this framework, biological traits refer to molecular or phenotypic measurements, such as gene expression levels, metabolite concentrations, or microbial abundances; whose pairwise relationships form the network. Modifying factors, by contrast, represent potential modulators of these relationships, including germline variants (e.g., SNPs), somatic mutations, copy number alterations, or other high-dimensional features. Since it is computationally prohibitive to analyze all possible triplets, GFBioNet employs a two-step procedure: first, it estimates a baseline network without considering modifiers, and second, it tests only those trait pairs (*i, j*) identified in the baseline network for evidence of modification by factor *t*. Under this framework, we provide a computationally efficient analytic approach to explicitly control false discovery rate (FDR).

Related methods include differential GGM (dGGM) analyses^16^, which compare networks across predefined conditions such as cases versus controls. A more recent penalized regression method, GMMReg^21^, estimates subject-specific networks, but relies on cross-validation (CV) and Bayesian Information Criterion (BIC) for tuning parameter selection, resulting in very high FDR. In contrast, GFBioNet appropriately controls FDR.

GFBioNet has broad applications in genomic analyses and network biology. To demonstrate its versatility, we applied GFBioNet to several datasets and contexts. First, we analyzed gut microbiome taxonomic data from shotgun metagenomic sequencing together with targeted metabolomic profiles quantified by capillary electrophoresis–time-of-flight mass spectrometry (CE-TOFMS) to reveal microbial modulation of metabolite networks^22^, providing insights into the metabolic interplay within the microbiome. Second, we examined host genetic influences on oral microbiome networks using 16S rRNA gene sequencing data from the Prostate, Lung, Colorectal, and Ovarian (PLCO)^23^ Cancer Screening Trial. Third, we explored somatic mutational events, including somatic copy number alterations (SCNAs) and somatic nucleotide variants (SNVs) in driver genes, linked to gene expression networks in The Cancer Genome Atlas (TCGA)^24^, shedding light on the regulatory disruptions associated with cancer. By expanding the scope of traditional GGM-based approaches to identify factors that modify direct pairwise relationships among traits, GFBioNet highlights the potential for uncovering novel biological insights and advancing our understanding of complex molecular networks.

## Results

### Overview of Methods

Given data on *M* biological traits, denoted as *x*_*ni*_ for subject *n* and trait *i*, and *T* genomic factors, denoted as *g*_*nt*_ for factor *t*, for *N* subjects, our goal is to identify triplets (*i, j, t*) with 1 ≤ *i* < *j* ≤ *M* and 1 ≤ *t* ≤ *T* such that the subject-specific PCC 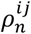 between traits *i* and *j* depends on the genomic factor *g*_*nt*_. We model this dependence as a linear function of all genomic factors

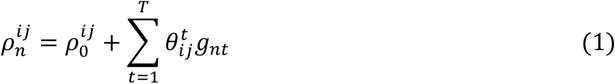

and the task reduces to identify nonzero coefficients 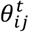. Although one could approach this problem by estimating a sparse precision matrix (**Methods**), that framework is both analytically and computationally challenging. Instead, identifying nonzero 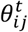 in equation (1) is equivalent to identifying nonzero 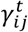, the coefficient measuring how factor *t* modifies the relationship between traits *i* and *j*, in the following regression model for *i* = 1, ⋯, *M*:

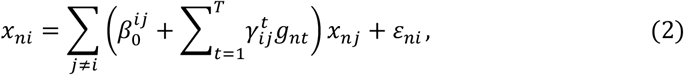

which is motivated by the nodewise regression approach for estimating GGM that leverages the relationship between regression coefficients and PCCs, which is similar to the method for identifying subject-specific GGM. Because the total number of variables far exceeds the sample size, we estimate the parameters using regularized regression with a penalty term 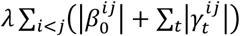. However, this framework requires estimating approximately *TM*^2^/2 parameters, making it computationally prohibitive and statistically underpowered due to the substantial multiple testing burden.

We propose a two-stage approach to address this problem (**Figure 1c**). In the first stage, we apply penalized nodewise regression to the *M* traits to estimate the baseline network. For each node (trait) *i*, let *ne*_*i*_ denote its neighborhood, i.e., the set of traits *j* such that 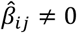 in the estimated baseline network. In the second stage, we restrict the search for non-zero nonzero 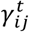 to pairs (*i, j*) with *j* ∈ *ne*_*i*_, by fitting:

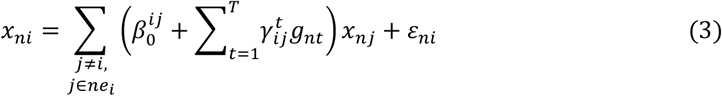

with penalty

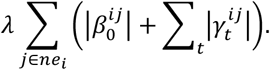

In both steps, it is crucial to select the tuning parameter *λ* to control the type-I error rate. CV and BIC have been used for estimating GGMs^2,3,25^. Recently, GMMreg^21^ was proposed for estimating personalized GGM; however, as we demonstrate, GMMreg produces very dense networks with unacceptably high false positive rates. We evaluated five methods for controlling FDR: CV, BIC, an analytic approximation, naïve permutation, and residual permutation (**Methods**).

The analytic approximation is derived from the penalized regression framework (LASSO or Elastic Net) under the sparsity assumption that only a small fraction of predictors represents true signals. Specifically, we establish a direct relationship between the tuning parameter (*λ*) and the expected number of false selections in penalized regression, allowing FDR to be approximated without computationally intensive permutations. This approximation leverages the asymptotic normality of the coordinate-descent updates and adjusts for predictor correlations through bias correction. As a result, GFBioNet can efficiently identify *λ* values that achieve the desired FDR level in high-dimensional settings while maintaining statistical power. A detailed derivation and justification of this approach is provided in the Supplementary Materials.

Although general post-selection inference methods can be used for FDR control^26,27^, they require estimating the precision matrix of all *T* × *M* predictors, which is a much more difficult problem in our setting. In contrast, permutation-based methods provide model-free calibration but vary in robustness^28^. Naïve permutation disrupts both signal and noise, inflating error rates when predictors are correlated, whereas residual permutation conditions on the fitted model and permutes only the noise, improving accuracy but requiring separate permutation for each candidate *λ*, making it computationally intensive.

Finally, before network estimation, each biological trait is adjusted for key covariates to remove confounding and improve statistical power. The residuals from these adjustments are used as input traits for network analysis. In addition, traits are quantile-normalized, and each genomic factor is standardized to unit variance prior to analysis.

### Evaluation of False Discovery Rate and Sensitivity in Simulations

We performed extensive simulations to investigate the performance of the algorithms, focusing on FDR and sensitivity. In each simulation, we modeled biological traits as continuous variables representing gene expression levels and genomic factors as SNPs acting as potential modulators of trait–trait relationships.

We generated a baseline network consisting of *M* = 100 nodes (biological traits) and 40 edges, where each edge represents a direct association (PCC) between the expression levels of two genes. The network included 10 hubs, defined as nodes connected to four other nodes (4 edges × 10 hubs). We then randomly selected 10 edges to be modulated by a SNP with effect size *γ* (**Figure 2a**) under an additive effect model (**Methods**). **Figure 2b** illustrates the relationship between *γ* and the corresponding PCC. Across simulations, we varied three parameters: sample size (*N*), effect size (*γ*), and the correlation among SNPs. We performed 50 simulations for each setting.

**Figure 2.**
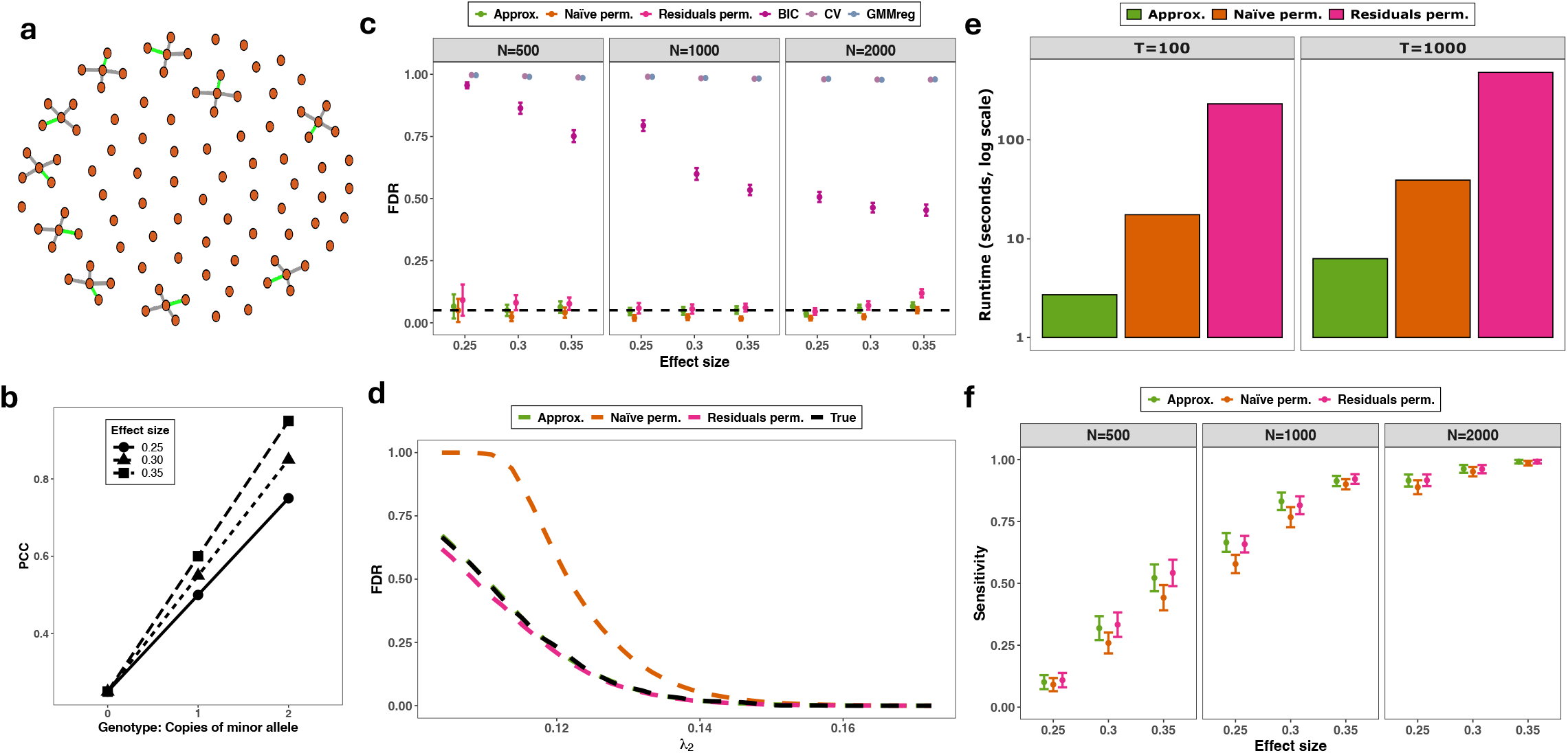
FDR and sensitivity in simulations. (a) Baseline network with 40 edges (4 edges × 10 hubs), 10 of which (green) were modulated by a SNP. (b) Relationship between effect size and PCC across three genotype groups. (b) Empirical FDR of GMMreg and GFBioNet under five tuning-parameter selection methods (50 simulations). (d) FDR curves for GFBioNet as a function of λ_2_ in the second-stage analysis. Relative to the true FDR (black dashed line), naïve permutation is conservative, residual permutation slightly liberal, and analytic approximation controls FDR well. (e) Computational time for GFBioNet using analytic approximation, naïve permutation, and residual permutation. (f) Sensitivity of GFBioNet under the three tuning-parameter selection methods.

We first evaluated FDR using five methods (analytic approximation, naïve and residuals permutation, CV, and BIC) for choosing *λ* based on a LASSO penalty. We also compared these with GMMreg (one-step procedure using CV to select *λ*). For naïve and residual permutations, we used 20 permutations to obtain stable FDR estimates.

Procedures using CV or BIC yielded unacceptably high FDR (**Figure 2c**). GFBioNet with naïve permutations was generally conservative (**Figures 2c** and **2d**), whereas GFBioNet with residual permutations was slightly liberal with small sample sizes (**Figure 2c**). Encouragingly, GFBioNet based on analytic approximations controls FDR well across all simulations (**Figures 2c** and **2d**) and was computationally much more efficient compared to permutation-based methods, particularly residual permutation (**Figure 2e**). We also performed simulations using Elastic Net penalty in our implementation of GFBioNet and obtained similar results (**Supplementary Figure 1, 2**).

Next, we investigated the computation cost of GFBioNet based three methods (analytic approximation and the two permutation methods) for choosing tuning parameters. Across all settings, the analytic approximation was consistently fastest (**Figure 2e**). For example, with *T* = 1000, the analytic approximation was 6x faster than naïve permutation and 75x faster than residual permutation. These results highlight the substantial computational advantage of the analytic approach, especially in high-dimensional settings.

Next, we evaluated the sensitivity of the GFBioNet using three methods. As expected, sensitivity increased with larger sample sizes and stronger effect sizes. When the effect size was reasonably large, GFBioNet exhibited robust statistical power (**Figure 2f**), particularly in scenarios where the number of traits exceeded the sample size, demonstrating its effectiveness in high-dimensional and sparse settings. Notably, GFBioNet with naïve permutations had lower sensitivity because it selected a more conservative *λ*.

In summary, our simulations showed that methods based on CV or BIC had unacceptably high FDR, whereas GFBioNet with the analytic approximation controls FDR effectively across all simulation settings while being more computationally efficient than permutation-based methods. Therefore, in subsequent real-data applications, we employed GFBioNet with the analytic approximation method for tuning parameter selection.

### GFBioNet identifies gut microbial taxa that modulate the fecal metabolite network

Recently, fecal metagenomic and metabolomic features were found to differ significantly among healthy controls, early-stage, and advanced-stage colorectal cancer (CRC) in a Japanese cohort^22^. In this study, 347 subjects with paired fecal metagenomics and fecal metabolomics data provided a unique opportunity to explore interactions between the gut microbiome and metabolites. While associations between metagenomic features and metabolites is often studied within a multi-omics analytical framework^29^, our aim here was to identify microbial taxa that modulate edges in the metabolite network.

We applied GFBioNet to the integrative dataset of 347 CRC patients with *M* = 121 metabolites and *T* = 224 microbial genera present in at least 70% of samples. For genus-level abundances, we performed the centered log-ratio (CLR) transformation to account for the compositional nature of the data. Analyses were adjusted for age, sex, body mass index (BMI), Brinkman Index (smoking history), alcohol consumption, and CRC stage.

At FDR ≤ 5%, the baseline metabolite network comprised 1,085 edges (**Figure 3**a, **Supplementary Table 1**), involving the 121 distinct metabolites. Among them, N-Acetyl-L-glutamic acid and S-Adenosylmethionine had the highest connectivity in the network, showing 31 and 28 direct associations with other metabolites, respectively.

**Figure 3.**
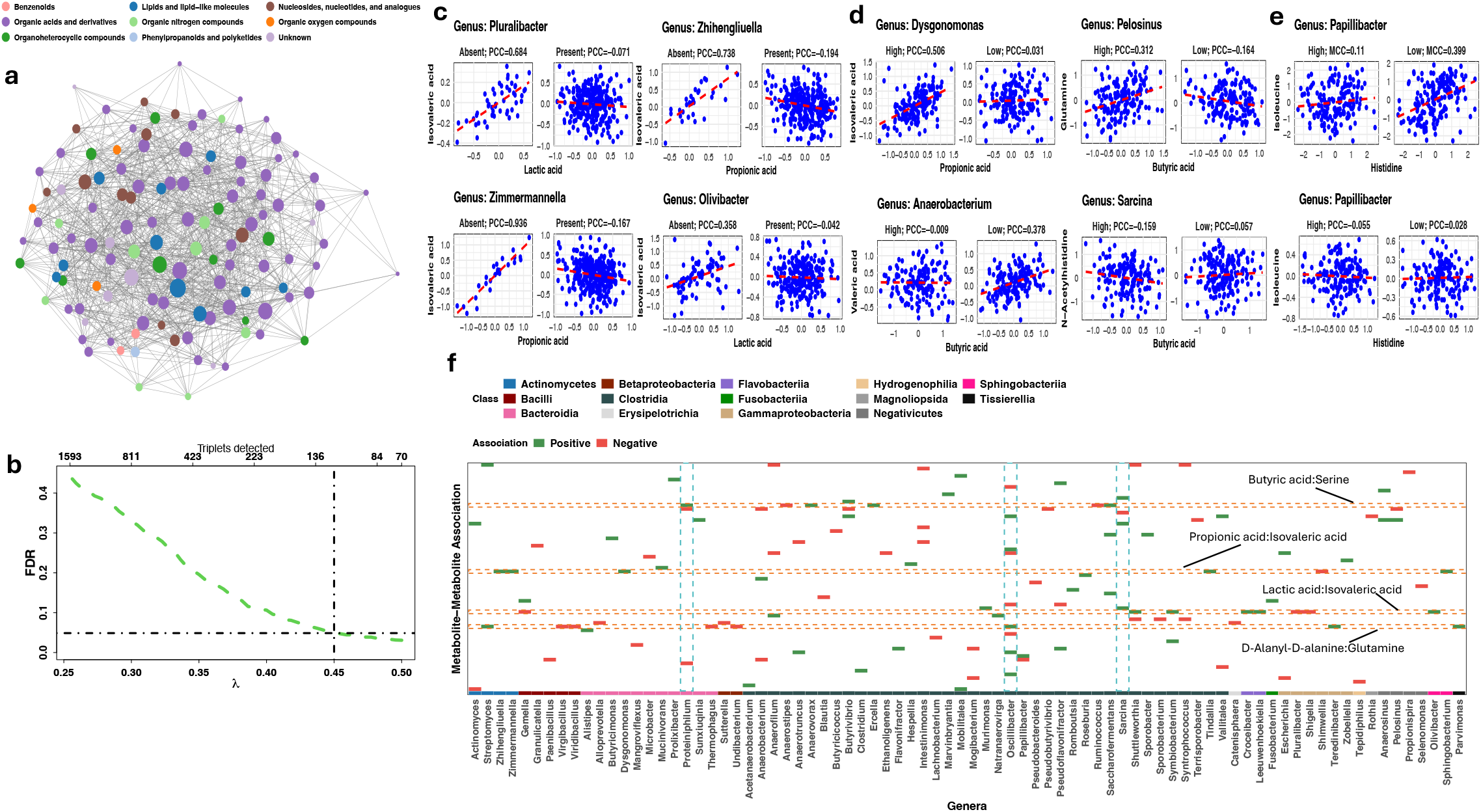
Gut microbial taxa associated with metabolite network structure in fecal samples from CRC patients in a Japanese cohort. (a) Baseline metabolite network (FDR 5%) with 1,085 metabolite pairs. Nodes represent metabolites, colored by class. (b) Relationship between FDR and number of detected signals as a function of tuning parameter λ; dashed line marks FDR = 0.05. (c) PCC between metabolites, stratified by presence/absence of genera. (d) PCC between metabolites, stratified by high/low CLR-transformed abundance of a specific genus. (e) Example: Papillibacter modifies the marginal but not partial correlation between isoleucine and histidine. (f) Summary of 120 metabolite–metabolite–genus triplets (FDR 5%). Columns = metabolite pairs; rows = genera. Green (red) boxes = positive (negative) PCC modification. Orange dotted horizontal lines = metabolite pairs modulated by ≥7 genera. Green dotted vertical lines = genera modulating ≥5 metabolite pairs.

GFBioNet identified (FDR ≤ 5%) 120 metabolite-metabolite-genus triplets, including 62 metabolites pairs and 80 unique genera (**Figure 3b, Supplementary Table 2**). These signals indicate microbial genera that modulate the PCC between a pair of metabolites. To illustrate that these results are supported by the data, we examined metabolite–metabolite correlations stratified by genus presence/absence or by dichotomized abundance (low: below median; high: above median). For example, the partial correlation between isovaleric acid and propionic acid was strongly positive when the genus *Pluralibacter* was absent (PCC=0.684) but was negative when the genus was present (PCC=-0.071; **Figure 3c**). In another example, *Dysgonomonas* abundance altered this association from PCC = 0.506 (high abundance) to PCC = 0.031 (low abundance; **Figure 3D**). Together, these examples suggest that genus-level presence or abundance can significantly influence metabolite–metabolite interactions.

In addition, we present an example where a signal identified by marginal correlation coefficient (MCC) analysis disappears when using PCC (**Figure 3e**). For two metabolites, isoleucine and histidine, the genus *Papillibacter* significantly modified their marginal association (MCC = 0.399 vs. 0.11 for high vs. low abundance). However, PCC analysis in the GGM framework showed no evidence of modulation (PCC = –0.055 vs. 0.028). This highlights the importance of distinguishing direct associations from indirect correlations using PCCs when interpreting microbial influences on metabolite networks. Direct associations identified through partial correlations are more likely to reflect metabolite relationships mediated by specific microbial processes, whereas indirect correlations (measured by MCC) may arise from the confounding of other metabolites.

Significant results are summarized in **Figure 3f**. We identified three genera, *Oscillibacter, Sarcina and Proteiniphilum*, as modulators of multiple metabolite pair associations. Among them, *Oscillibacter* emerged as a key modulator, influencing nine metabolite pairs. The modulation of butyric acid, a short-chain fatty acid with anti-inflammatory and tumor-suppressive properties, highlights *Oscillibacter*’s role in gut immune modulation and epithelial homeostasis^30^. Similarly, the Methionine–Methionine sulfoxide interaction suggests *Oscillibacter* may influence oxidative stress and sulfur metabolism, processes closely linked to CRC development^31^. Additionally, the Agmatine–Spermidine association points to its involvement in polyamine pathways that govern cell growth and apoptosis^32^. Collectively, these findings suggest that *Oscillibacter* may affect multiple metabolic pathways simultaneously, potentially through shared enzyme activity, metabolite exchange, or redox balance, thereby reshaping the topology of the metabolite network and exerting pathway-level effects relevant to CRC progression^33^.

Further, several metabolite pairs modulated by multiple genera, including D-Alanyl-D-alanine:Glutamine, Lactic acid:Isovaleric acid, Butyric acid:Serine, and Propionic acid:Isovaleric acid. For example, we identified eight genera (including *Parvimonas* and *Oscillibacter*) as modulators of the association between D-Alanyl-D-alanine and Glutamine (**Figure 3f, Supplementary Table 3**). D-Alanyl-D-alanine is essential for bacterial cell wall synthesis, while Glutamine plays a key role in gut epithelial repair, immune function, and nitrogen metabolism. The modulation of this interaction by multiple genera suggests microbial contributions to host-microbe metabolic interactions critical for maintaining gut health. Altered microbial composition involving genera such as *Parvimonas* and *Oscillibacter*, previously linked to CRC, may exacerbate inflammation and metabolic disruptions, potentially contributing to CRC progression^30,34^. This finding underscores the complex interplay between microbiota and host metabolic pathways in CRC development.

### Identification of Host Genetic Variants Modulating the Oral Microbiome Network

Microbes form complex, structured communities characterized by diverse interactions. The relationships among bacteria (cooperative, competitive, or neutral) play a critical role in shaping microbial composition and function. Host genetics may influence microbial communities across body sites, as demonstrated through heritability analysis^35^ and genome-wide association studies^36^ using taxa abundance or alpha diversity as traits. However, the role of host genetics in shaping microbial interactions, such as conditional associations or pairwise associations^6,37^, remains unexplored.

Here, we applied GFBioNet to identify host genetic variants associated with the oral microbiome network in 1,356 subjects from the PLCO cohort^23^ (**Methods**). The biological traits analyzed were CLR-transformed abundances of 76 genera, present in at least 20% of subjects. After quality control and filtering (**Methods**), 470,66*s* common SNPs remained for analysis. Analyses were adjusted for relevant demographic characteristics (**Methods**) and principal component scores derived from host genotypes to account for population stratification. At an FDR threshold of 5%, we identified a baseline network comprising 666 edges among the 76 genera (**Figure 4a**).

**Figure 4.**
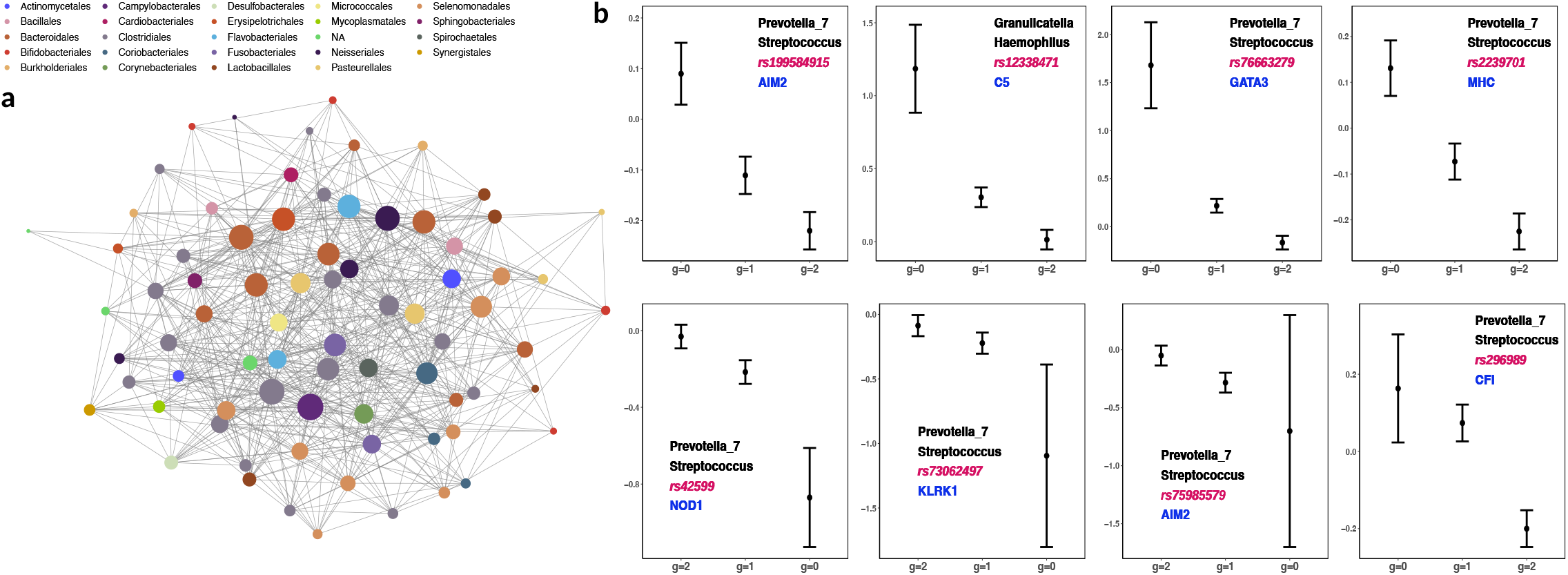
Host genetic variants modify genus-level interactions in the oral microbiome in the PLCO study. (a) Baseline network of 76 genera in the oral microbiome. Node colors denote taxonomic class; node size reflects connectivity. (b) Eight genetic signals were identified (FDR 10%), focusing on genetic variants in the MHC region or immune-related loci. Panels show Fisher-transformed PCCs across three genotype groups (0, 1, 2 copies of the minor allele); error bars indicate standard errors, driven largely by group sample sizes. In each panel, the modulated genus pair is annotated in black, the corresponding single-nucleotide polymorphism (SNP) in red, and the nearby gene in blue.

The initial analysis included all 470,669 SNPs but did not identify significant signals at FDR 10%. Previous studies have demonstrated that SNPs in the major histocompatibility complex (MHC) region and immune-related genes are likely associated with microbiome abundance and diversity^38-40^; therefore, we hypothesized that these variants might also influence microbiome interactions. To test this, we restricted our analysis to 13,309 SNPs located within the MHC region or immune genes (Methods). This refined analysis identified eight genus-genus-SNP triplets at FDR ≤ 10% (**Figure 4b**), involving two genus-genus interactions: *Prevotella_7* – *Streptococcus* and *Granulicatella* – *Haemophilus*.

The interaction between *Prevotella_7* and *Streptococcus* was modulated by seven SNPs across six immune-related loci. These variants were not associated with the relative abundance of either genus (data not shown), suggesting that host immune genetics influence the interaction between these microbial taxa rather than their individual abundances. The genes nearby the identified SNPs, including *TAP1/TAP2* (antigen presentation), *NOD1* (immune signaling), *KLRK1/KLRC1/KLRC2* (*NK* cell activation), *AIM2* (inflammasome regulation), and *CFI* (complement activity), play key roles in mucosal immunity and may shape *Prevotella_7– Streptococcus* interactions^41,42^. The *Prevotella* genus is associated with high-fiber diets, lower mucosal inflammation, and periodontitis^41^, while *Streptococcus* includes both commensal and pathogenic species affecting oral health^43^. The observed host genetic effects suggest that immune modulation influences microbial interaction, potentially affecting disease susceptibility.

In addition, the interaction between *Granulicatella* and *Haemophilus* was modulated by a SNP within the immune gene *C5*, a key component of the complement cascade involved in immune defense and inflammation^44^. *Granulicatella* is an oral commensal bacterium linked to endocarditis^45^, while *Haemophilus* includes species associated with oral and respiratory infections^46^. Given that these genes are linked to autoimmune diseases, the interplay between host genetics, microbial interactions, and inflammatory processes warrants further investigation^47^.

### Identification of Somatic Mutations Modulating Gene Expression Networks in LUAD

Driver genes have been discovered for many tumors in sequencing studies by comparing the frequencies of nonsynonymous mutations to background mutations^46^. However, the potential functional mechanisms through which these mutations influence tumor initiation and progression remain unclear for many driver genes. Here, we applied GFBioNet to explore how somatic mutations modulate gene expression networks in LUAD, by leveraging data from TCGA^24^.

The first-stage analysis included 4,872 genes with the most variable expression levels from 279 stage I/II LUAD patients of European ancestry (**Methods**) and estimated a baseline network consisting of 80,016 edges, where each edge represents a direct association (PCC) between the expression levels of two genes. The second-stage analysis focused on the estimated baseline network and 16 LUAD driver genes (**Methods**) with binary mutation data^24^. By controlling FDR at 1%, GFBioNet identified 109 expression-expression-mutation triplets involving 13 driver genes. Among these, most somatic mutations increased PCCs (100 out of 109 triplets), while *TP53* mutations consistently decreased PCCs across four pairs of gene expression levels. Mutations in *KRAS, KEAP1*, and *STK11* demonstrated the highest modulatory impact, influencing 35, 23, and 13 gene pairs (**Figure 5a**), respectively.

**Figure 5.**
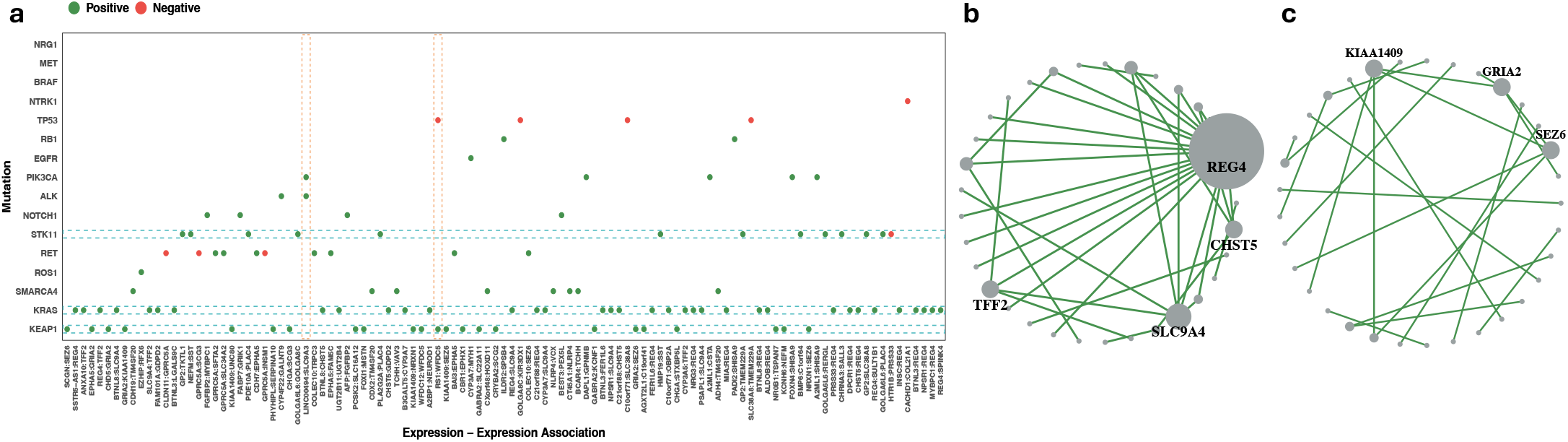
Somatic driver gene mutations alter gene expression networks in LUAD in TCGA. (a) Overall impact: GFBioNet identified 109 expression-expression–mutation triplets in which driver mutations modified partial correlations (PCCs) between pairs of gene expression levels (FDR ≤ 0.01). (b) *KRAS*-associated dependencies: PCCs between gene expression levels were significantly elevated in *KRAS*-mutant tumors. Nodes represent individual genes, and edges indicate direct associations whose PCCs were modified by *KRAS* mutation status. Node size corresponds to the number of *KRAS*-modified associations per gene. (c) *KEAP1*-associated dependencies: PCCs between gene expression levels were significantly increased in *KEAP1*-mutant tumors.

Of the 35 gene pairs with increased PCC between their expression levels due to *KRAS* mutations (**Figure 5b**), 18 involved *REG4*, which enhances cancer stem cell traits in KRAS-mutant colorectal cancer^48^ and marks a LUAD subtype with low *TTF*-*1* expression^49^, consistent with poorly differentiated tumors. Although *REG4* is associated with poor prognosis in multiple cancers^50^, its role in most LUAD remains unclear. The enrichment of *REG4*-related edges in *KRAS*-mutant tumors suggests activation of a mucin-producing transcriptional program, reminiscent of an intestinal-like phenotype observed in TTF-1–low LUAD^49^. The interaction between expression levels of gene pairs such as *REG4–TFF2, REG4–SPINK4*, and *SLC9A4– REG4* suggest roles in mucosal defense and epithelial regeneration, while *REG4–BTNL8* and *REG4–BTNL3* imply immune modulation via BTNL proteins. Together, these findings support the hypothesis that *KRAS* mutations promote mucosal protection, epithelial repair, and immune modulation, warranting further investigation into *REG4*’s molecular role and therapeutic potential.

*KEAP1* modulates the *NRF2* pathway^51^, which manages oxidative stress responses^52^. Mutations in *KEAP1* can lead to uncontrolled *NRF2* activity, promoting tumor progression and poor prognosis^53^. In our analysis, *KEAP1* mutations increased PCCs between the expression levels of 24 gene pairs (**Figure 5c**), particularly involving *GRIA2* and *SEZ6*, each participating in four pairs. This observation may indicate a shift in cellular processes within LUAD tumors, suggesting that *KEAP1* mutations could influence pathways beyond oxidative stress response, potentially affecting cellular communication and signaling mechanisms. Better understanding could provide insights into novel therapeutic targets for *KEAP1*-mutant LUAD.

### Identification of Somatic Copy Number Alterations (SCNAs) Modifying Gene Expression Networks in LUAD

Finally, we explored the impact of SCNAs on gene expression networks in LUAD using the same dataset, which included the baseline network of 80,016 edges described in the previous section. The genome was partitioned into one-megabase segments, with SCNA measured as the average intensity within each segment. After correlation pruning and filtering out low-variance segments (**Methods**), 283 SCNA segments remained for analysis. We considered SCNA as a continuous variable and aimed to detect its trend effect on the PCCs between gene expression levels inferred from the gene expression network. To account for potential confounding factors, the analysis was adjusted for age, sex, tumor stage, PEER factors^54^, and tumor purity^55^.

At a 1% FDR threshold, GFBioNet identified 3,188 expression-expression-SCNA triplets were (**Figure 6a**), corresponding to 1,631 unique edges. Here, an edge represents a direct association (PCC) between the expression levels of a pair of genes, and an edge is considered modified when the strength or direction of this PCC varies systematically with SCNA segment values. Most edges (1,143) were influenced by a single SCNA segment, while 164 edges were associated with four or more segments. Among the 283 SCNA segments analyzed, 281 were linked to at least one modified edge, with a median of nine edges per segment, highlighting the widespread influence of SCNAs on transcriptional relationships across the genome.

**Figure 6.**
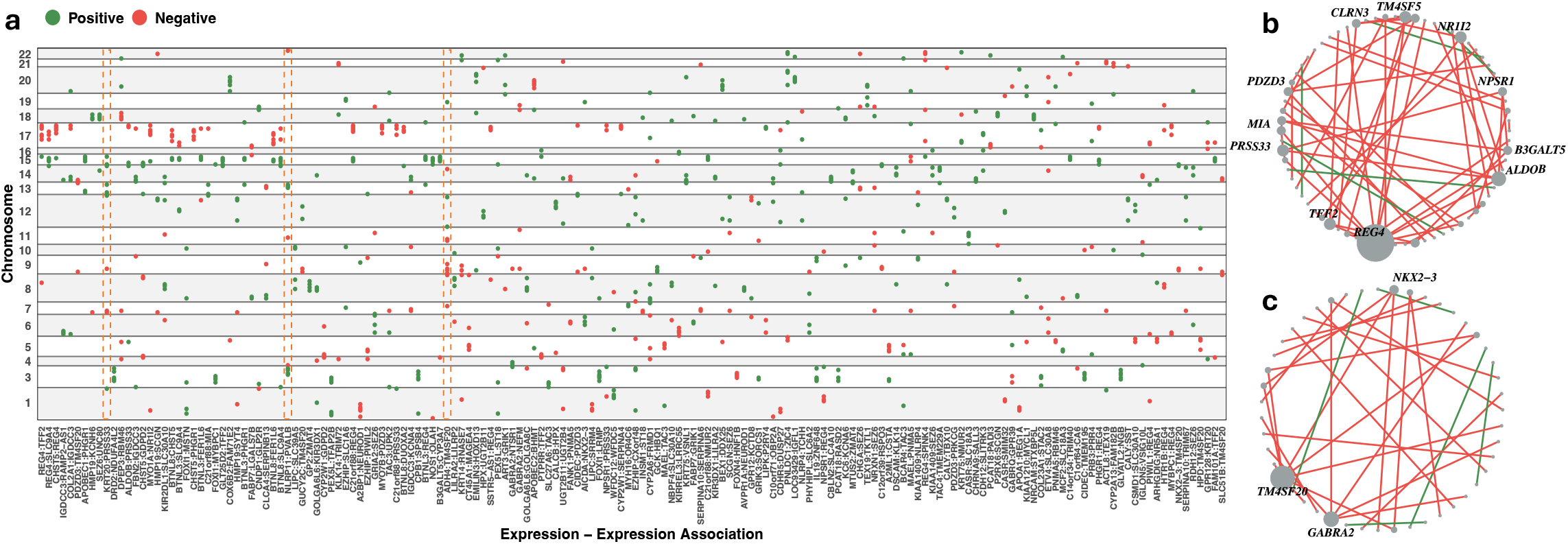
Somatic copy number alterations (SCNAs) influence gene expression dependencies in LUAD (TCGA). (a) Overall impact: Rows represent SCNA segments, and columns represent gene pairs. Shown are *KRT20:PRSS33; NLRP11:PVALB; ADH4:TM4SF20* whose partial correlations (PCCs) between gene expression levels were associated with ≥10 SCNA segments. Red and green dots denote positive and negative associations, respectively, between SCNA values and PCC strength. (b) *TP53*-associated dependencies: Gene pairs whose expression-level associations (PCCs) were most strongly influenced by the SCNA segment encompassing *TP53*. Node size reflects the number of gene expression dependencies affected by the SCNA segment. (c) *CDKN2A*-associated dependencies: Gene pairs whose expression-level associations were influenced by the SCNA segment containing *CDKN2A*.

The SCNA segment affecting the largest number of gene expression dependencies (51 edges) encompassed the *TP53* locus (**Figure 6b**), a key tumor suppressor implicated in most cancers. Deletions in this region led to increased PCCs for 48 of these gene pairs, reflecting its broad impact on gene-gene interactions and reinforcing *TP53*’s critical role in tumor biology. Similarly, SCNA segments harboring *CDKN2A* influenced 38 edges (**Figure 6c**), including 11 involving *TM4SF20*, a gene implicated in proliferation and oncogenic signaling. Additionally, 99 *REG4*-related edges were associated with SCNA segments, implicating *REG4*’s role in LUAD cell proliferation and differentiation, as observed in the previous analysis of driver gene mutations (**Figure 5b**). Together, these findings illustrate that SCNA events in key cancer-associated loci can reshape dependencies among gene expression levels, thereby reorganizing transcriptional networks relevant to LUAD cell proliferation and differentiation.

## Discussion

In summary, our work addresses a critical gap in current methodologies for integrating high-dimensional data with network-based approaches. Existing multi-omics analyses primarily focus on identifying factors associated with mean or variance shifts in individual traits. This node-centric perspective ignores the possibility that both genomic and non-genomic factors can act on the edges of biological networks, altering relationships between traits and reshaping network structure. By developing a robust statistical framework, we enable the identification of factors that modulate direct trait-to-trait relationships, moving beyond conventional network analyses of multi-omics data or differential network analyses. Our approach offers notable advantages while maintaining computational feasibility, including its flexibility in accommodating diverse data types and its ability to explicitly control FDR. These features ensure broad applicability across various biological and molecular contexts. The two-stage strategy, which restricts the search for modulators to edges in the estimated baseline network, substantially reduces computational and multiple-testing burdens while increasing the power to detect meaningful associations. This methodological advancement paves the way for future research into the genomic modulation of biological networks.

Although our applications focus on genomic factors (e.g., genetic variants, somatic mutations, SCNAs) as modulators of direct trait–trait relationships, the framework is not limited to genomics. Any well-measured non-genomic factors, such as environmental exposures, drug treatments, clinical phenotypes, or experimental conditions, can be incorporated as an external variable influencing network edges. Both continuous and categorical modulators are supported, and multiple modulators can be considered jointly. Thus, GFBioNet offers a flexible tool for studying how diverse biological or environmental factors modulate pairwise dependencies among traits.

A persistent challenge in network analyses using GGM is the selection of appropriate tuning parameters to control type-I error rates. Frequently used approaches such as the CV and BIC for model selection often yield excessively dense networks with high FDR. To address this, we developed and evaluated three methods to control FDR. Extensive simulations showed that the analytic approximation provides the most reliable FDR control while being substantially more computationally efficient than the two permutation-based methods. While the regression framework of GFBioNet is similar to that of GMMreg, GMMreg is a one-step procedure that cannot handle high-dimensional data efficiently and results in unacceptably high false positive rates.

As illustrated throughout this study, GFBioNet can be broadly applied to key problems in multi-omics analyses. The methodological framework can be extended to tackle additional analytical challenges. First, the sensitivity of network analyses often benefits from explicitly modeling “hubs”, traits that maintain direct associations with multiple other traits^17,56,57^. Extending this concept to our framework, detecting factors that influence multiple trait–trait dependencies could enhance sensitivity. Indeed, GFBioNet identified multiple hotspot-modifying factors, e.g., the SCNA segments encompassing *TP53* and *CDKN2A* (**Figure 6**). Extending GFBioNet to specifically detect such hotspot-modifying factors could further improve power and interpretability. Second, the method holds potential for application in single-cell RNA sequencing studies, where genetic variants or environmental exposures may modulate gene expression networks across different cell types. Aggregating evidence across multiple cell types could enhance the statistical power for detecting modulators of gene expression network in single-cell data, opening new avenues for studying cell-type-specific gene modulatory mechanisms. Finally, while our two-stage strategy effectively reduces computational complexity and improves statistical power, it may miss true signals for trait pairs that are not present or have different directions of association in the baseline network. A potential future extension could involve adapting the stratified FDR procedure to allow distinct tuning parameters for trait pairs within and outside the baseline network. This refinement could balance power and false positive control while retaining computational efficiency.

Overall, GFBioNet provides a useful tool for improving our understanding of how high-dimensional factors (genetic or otherwise) modulate the conditional relationships among biological traits. By addressing key methodological challenges in statistical genomics and network biology, the framework has broad applicability across complex biological systems and may inspire future methodological innovations.

## Methods

### A Brief Overview of Gaussian Graphic Models

GGMs are widely used to explore association relationships among many variables. Conceptually, GGMs aim to identify pairs of variables with non-zero PCC, which indicate a direct dependency between variables after accounting for the effects of all other variables. This contrasts with MCC-based analysis, which measures indirect relationships between variables. Let (*x*_*n*1_, …, *x*_*nM*_) be a vector for subject *n* drawn from a multivariate normal distribution *MVN*(0, *Σ*), where *Σ* denotes the covariance matrix. Define *Ω* = *Σ*^-1^ as the precision matrix. Lauritzen (1996)^1^ showed that the PCC *ρ*^*ij*^ between *i* and *j* can be expressed as

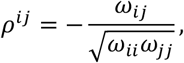

where *ω*_*ij*_ is the (*i, j*)-th entry of the precision matrix *Ω*.

Methods for estimating GGM can be grouped into three main categories. The first approach estimates a sparse precision matrix *Ω* by 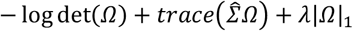, where 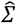 is the empirical covariance matrix^2^. Efficient algorithms, such as graphical LASSO (gLasso)^3^, have been developed. The second approach uses regularized nodewise regression^11^, leveraging the relationship between regression coefficients and PCCs. For variable *i* (node *i*), a linear regression model is constructed as follows:

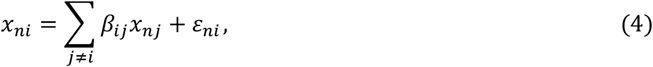

Where 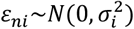. It has been shown that *β*_*ij*_ = −*ω*_*ij*_/*ω*_*ii*_ and *β*_*ji*_ = −*ω*_*ji*_/*ω*_*jj*_; thus, identifying non-zero PCC is thus equivalent to identifying non-zero *ω*_*ij*_, or equivalently, pairs where *β*_*ij*_ ≠ 0 and *β*_*ji*_ ≠ 0^1^. Given that the number of variables often far exceeds the number of observations, Meinshausen and Bühlmann proposed estimating regression models using penalized regression techniques, such as LASSO^58^ or Elastic Net^59^.

The third type of method consists of limited-order partial correlation methods. When calculating a PCC, these methods adjust for a subset of other traits rather than performing a full adjustment^12-14^. As a result, the derived PCC falls between the true PCC and MCC, and the resulting network lies between a GGM and a marginal association network.

### Identifying High-Dimensional Genomic Factors Associated with Biological Networks

We now address the problem of identifying high-dimensional genomic factors associated with biological networks represented as a GGM. Specifically, given data on *M* traits and *T* genomic factors for *N* subjects, we aim to identify triplets (*i, j, t*) for 1 ≤ *i* < *j* ≤ *M* and 1 ≤ *t* ≤ *T*, such that the subject-specific PCC 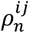 depends on the genomic factor *g*_*nt*_. We assume that trait values *x*_*ni*_ and genomic factors *g*_*nt*_ are normalized to have mean zero and unit variance across subjects. We model the subject-specific PCC as a linear model of all *T* genomic factors:

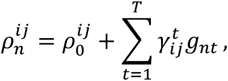

where 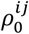 denotes PCC between *i* and *j* averaged across the cohort. Identifying genomic factors associated with a GGM is equivalent to identifying nonzero 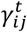. While we can approach the problem via the precision matrix, it is extremely challenging, particularly when the dataset includes thousands of traits and millions of genomic factors.

The alternative is to extend the nodewise regression approach. In equation (4), we model subject-specific coefficient 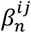 as:

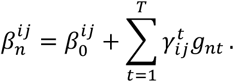

By introducing an index for subject *n* and plugging 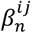 into the linear regression model (4) for node *i*, equation (4) reduces to

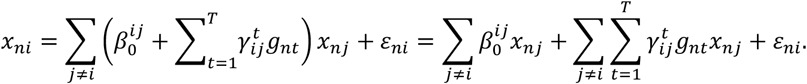

Thus, identifying the population-averaged GGM and genomic factors associated with GGM requires to identifying node pairs (*i, j*) with 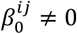 and identifying triplets (*i, j, t*) with 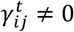, respectively, which is solved by the following penalized regression for each node *i* by minimizing

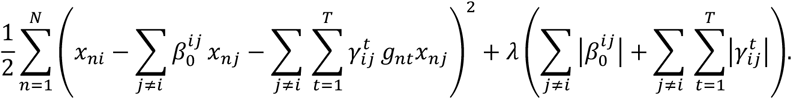

From the perspective of regression analysis, 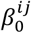 measures the main effect of *x*_*j*_ on *x*_*i*_ and 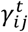 measures the interaction effect of *x*_*j*_ and genomic factor *g*_*nt*_.

This framework is similar to GGMreg^21^ for estimating subject-specific GGM; however, GGMreg uses CV or BIC to choose appropriate tuning parameter *λ* and has very high FDR based on our simulations, suggesting that most of the identified signals are false positives. We will develop computationally efficient methods to explicitly control FDR.

### Two-stage Procedure for Identifying Genomic Factors Associated with Biological Networks

There are *M*(*M* − 1)/2 main effects and *M*(*M* − 1)T/2 interactions to be estimated, presenting two major challenges when *M* and *T* are large. For example, in the application of identifying common host genetic variants (*T* ≈ 7 million) associated with the microbiome network (*M* = ∼300 after filtering), there are approximately 4.5 × 10^4^ main effects and 3 × 10^11^ interactions, making computation prohibitive. Furthermore, the large number of interactions requires selecting a large *λ* value to control FDR, reducing statistical power. To address these challenges, we propose a two-stage procedure.

In the first stage, we construct the baseline network using nodewise penalized regression. For node *i*, we solve the penalized regression problem by minimizing:

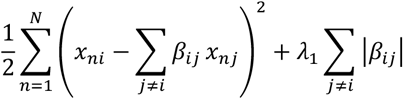

to derive the regularized estimate 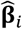. The neighborhood of node *i*, denoted as *ne*_*i*_ includes nodes with nonzero coefficients. We performed regularized nodewise regression for each of the nodes and denoted the resulting network as an undirected graph 𝒢 = (𝒱, ℰ), where 𝒱 denotes the set of vertices with non-empty neighborhood and ℰ is the adjacency matrix. We assume that the baseline network has *𝜈* = |𝒱| vertices and *d* = |ℰ| edges.

The second stage aims to identify a subset of *d* × *T* variant-edge interactions, where each interaction specifies a genomic factor influencing the association (edge) between two traits. By extending the neighborhood selection approach, we fit *v* penalized regression models to detect such interactions. For each *i* ∈ 𝒱 in the baseline network, we fit the penalized regression model

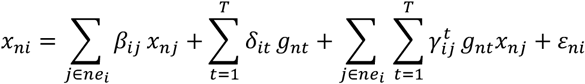

by minimizing

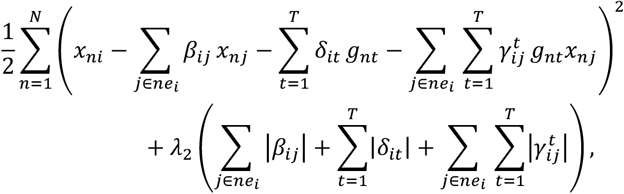

where *λ*_2_ > 0 is a tuning parameter.

For notational simplicity, we have ignored covariates; but the statistical framework can be easily modified to adjust for covariates, which include demographic characteristics, population stratification vectors measured by principal component vectors when investigating genetic variants, tumor purities when analyzing gene expressions and SCNA, probabilistic estimation of expression residuals (PEER) when analyzing gene expressions.

### Choosing Tuning Parameter to Control False Discovery Rate in Regularized Regression

Results in this section will be used to derive a procedure to control FDR in network analysis in the next section. Details are in Supplementary Materials. Consider the LASSO regression

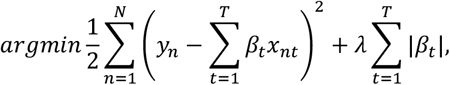

where *x*_*nt*_ and *y*_*n*_ are normalized to have mean zero and variance one. Let 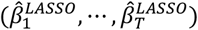 be the LASSO estimator given *λ*. Let 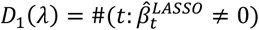 be the number non-zero estimates, and let

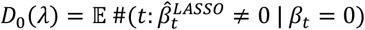

be the expected number of false positives. Then, FDR is estimated as *D*_0_(*λ*)/*D*_1_(*λ*) and we can find an appropriate *λ* to control FDR at an appropriate level. In Supplementary Materials, we develop the approximation:

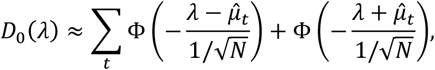

where Φ is the cumulative distribution function of 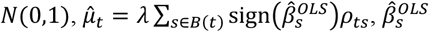, is the OLS estimator of the last step of coordinate descent algorithm, *ρ*_*st*_ = cor(*X*_*ns*_, *X*_*nt*_), and 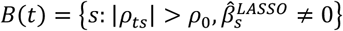. This approximation is derived under the assumption of sparseness, i.e., only a small fraction of predictors are causal. Implementing the algorithm is computationally efficient. For predictor *t*, we calculate 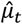 by only including predictors in *B*(t), which have non-zero LASSO estimate (just a few when *λ* is not small) and are correlated with *X*_*t*_.

For the Elastic Net regression 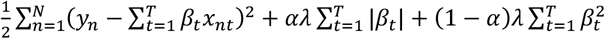, we used the similar approach to derive approximation

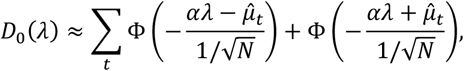

where

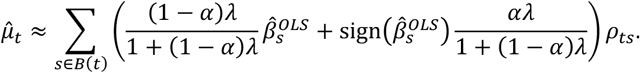

For the special case with *α* = 1/2, we have

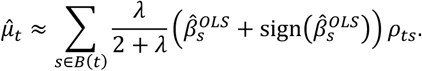

### Choosing Tuning Parameters to Control False Discovery Rate in GFBioNet

Our two-step procedure requires choosing tuning parameters *λ*_1_ and *λ*_2_, where *λ*_1_ determines the size of the baseline network and *λ*_2_ determines the number of signals of interest. Popular methods include CV^2^ and BIC^2,3,25^; however, both types of methods tend to have unacceptably high false positive rates, leading to very dense false network^60^. Liu et al. (2010) proposed a resampling-based approach and a new stability criterion that significantly reduced the false positive rate. While successfully reducing false positive rates, this approach does not explicitly control false positive rates at a desirable level. In addition, it is not straightforward to extend this approach to address the challenge in our problem. van de Geer, Bühlmann, Ritov, and Dezeure (2014) and Javanmard & Montanari (2014) proposed a statistical framework for post-selection inference for LASSO regression^26,27^. This framework provides debiased estimate and standard errors, which allows deriving p-values and controlling FDR. However, this approach requires estimating the sparse precision matrix of all predictors, which poses an even bigger challenge in our problem.

We aim to control FDR in the two-stage procedure. We choose *λ*_1_ to control FDR≤ *α*_1_ when building the baseline network in the first stage and choose *λ*_2_ to control FDR≤ *α*_2_ when identifying nonzero interactions 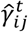 in the second stage. For stage one, let *D*_11_(*λ*_1_) denote the number of nonzero 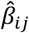 in the *M* regressions and *D*_10_(*λ*_1_) denote the expected number of false positives. For the second stage, we define *D*_21_(*λ*_2_) to be the number of non-zero interactions 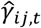 and *D*_20_(*λ*_2_) to be the expected number of false interactions. FDR is calculated as *FDR*(*λ*_1_) = *D*_10_(*λ*_1_) /*D*_11_(*λ*_1_) for the first stage and *FDR*(*λ*_2_) = *D*_20_(*λ*_2_) /*D*_21_(*λ*_2_) for the second stage. The key step of estimating FDR is to approximate *D*_10_(*λ*_1_) and *D*_20_(*λ*_2_) for the two stages, respectively. Here, we present three methods for estimating *D*_10_(*λ*_1_) and *D*_20_(*λ*_2_): analytic approximations, naïve permutations, and residual permutations.

### Analytic approximations

Analytic approximations to FDR depend on the approximation of *D*_0_(*λ*) in penalized regression, as we summarized in previous section. We illustrate the approach for selecting *λ*_2_ in the second stage analysis with *v* penalized regressions for trait *i* ∈ 𝒱:

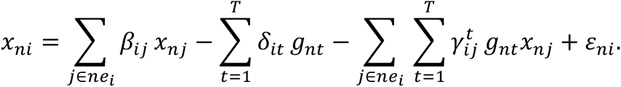

Given *λ*_2_, let 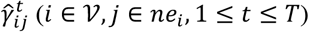 be the penalized estimate. Then,

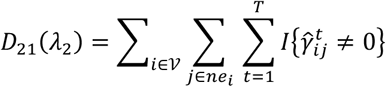

is the total number of detections with non-zero coefficients. Based on the approximation of *D*_0_ for LASSO regression in the previous section, the total number of expected false positives can be approximated as

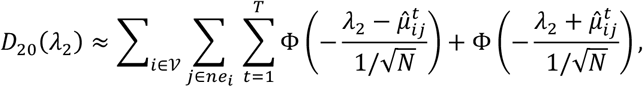

where 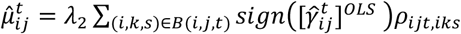. Here, *ρ*_*ijt,iks*_ = *cor*(*x*_*nj*_ *g*_*nt*_, *x*_*nk*_g_*ns*_) and 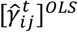 is the OLS estimate of 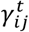 in the last round of iterations of the coordinate descent algorithm in the LASSO estimate. Similarly, we can approximate *D*_20_(*λ*_2_) when using Elastic Net regression to identify non-zero interactions.

### Naïve permutation

In the nodewise regression *x*_*ni*_ = ∑_*j*≠*i*_ *β*_*ij*_*x*_*nj*_ + *ε*_*ni*_ for stage one, we permute the response variables *x*_*ni*_ across subjects for *K* times. Then, we refit the regression each time to compute 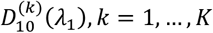. Finally, we estimate *D*_10_(*λ*_1_) as the average number of 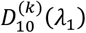. Similarly, for the regression in the second stag 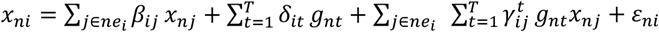, we permute the response variables *x*_*ni*_ across subjects for *K* times, refit the regression and estimate *D*_20_(*λ*_2_) as the average number of detected interactions.

### Residual permutation

For stage one, we first obtain the penalized estimate 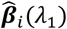 for node *i*, and calculate the residuals 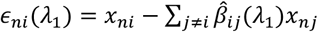. We then permute *ϵ*_*ni*_(*λ*_1_) and refit the regression *K* times, using the permuted residuals as responses. We estimate *D*_10_(*λ*_1_) by averaging the detections across *K* permutations. Similarly, we perform residual-based permutations to estimate *D*_20_(*λ*_2_) for stage two.

For naïve permutation, model parameters can be efficiently estimated across all candidate tuning values because the same *K* permutations are applied to compute the full solution path. In contrast, residual permutation depends on the residuals computed for each *λ*; therefore, a separate permutation is required for every tuning value. This makes residual permutation considerably more computationally intensive. It is thus important to narrow the search range for *λ* (see next section).

### Selecting Candidate Tuning Values for Residual Permutations

For the LASSO regression under the assumption of independent predictors, we derive an upper bound *λ*^*upper*^, corresponding to a family-wise error rate (FWER) of 0.05, and a lower bound *λ*^*lower*^, corresponding to the detection of at most *K* =10,000 signals. Although these bounds are derived assuming independence, they typically provide a reasonable range of *λ* values for FDR control.

We illustrate the method for stage one analysis. Because we have *M* regressions and each has (*M* − 1) coefficients to be estimated, the stage-one analysis requires estimating *M*(*M* − 1) parameters. To ensure a family-wise error rate of *α*_0_ = 0.05, we select *λ*_1_ such that

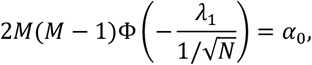

which leads to the upper bound of

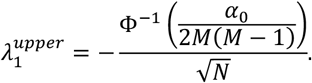

On the other hand, we assume that we detect *K* = 10,000 signals with tuning parameter *λ*_1_. Under this assumption, controlling *FDR* ≤ *α*_1_ leads to the lower bound

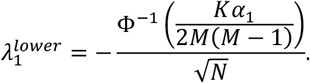

The upper and lower bounds for the stage-two analysis can be derived analogously under the same independence assumption.

### Host Genetics and Oral Microbiome Network

Oral microbiome and host genetic data from 1,362 subjects were obtained from PLCO. All subjects had no history of cancer and were of European ancestry, confirmed through both self-reported data and genotype-derived principal component scores. Following the lung cancer analysis in the same dataset^61^, our analyses were adjusted for demographic characteristics, including age, education level, BMI, alcohol consumption, and smoking behavior. Notably, smoking history was categorized using a 22-level variable, incorporating smoking status (current, former, never), cigarettes per day, and time since quitting.

The details for DNA extraction, PCR amplification, and sequencing from the buccal cell pellet was previously described. Briefly, DNA was extracted using the DSP DNA Virus Pathogen kit (Qiagen); PCR was performed using 16S rRNA gene V4 barcoded primers; and DNA was sequenced using the MiSeq (Illumina) with 2 × 250-bp paired-end sequencing. The sequencing data were processed using QIIME 2 version 2018.4 to generate amplicon sequence variants (ASVs) and create our feature table with DADA2. Taxonomy was assigned to ASVs using SILVA v132 classifier and the q2-feature-classifier plugin.

Genotype data were generated using the Illumina Array. Quality control (QC) was performed by excluding samples with a low call rate, non-European ancestry, discrepancies between self-reported and genotype-inferred sex, and cryptic relatedness. Additionally, SNPs with a call rate <0.95 or minor allele frequency (MAF <0.05) were removed. To reduce computational complexity and redundancy, we applied linkage disequilibrium (LD)-based pruning using PLINK, with a 500 kb window and an R^2^ threshold of 0.8.

### Lung Adenocarcinoma: Data Acquisition, Preprocessing, and Normalization

We downloaded processed meta data and genomic data from TCGA at https://www.cbioportal.org/datasets, including gene expression, SCNA, and somatic mutational data. To avoid heterogenicity, we only kept stage I/II LUAD patients of European ancestry.

#### Preprocessing and filtering of gene expression data

We chose the 5000 genes for analysis with the maximum variance across subjects. Because GGM assumes a multivariate normal distribution, we performed quantile normalization for each gene so that the gene expression levels roughly followed a normal distribution. To avoid the complexity of highly correlated genes, we calculated pairwise correlation between gene expression levels. For any pairs of genes with a correlation coefficient *R* with *R*^2^ ≥ 0.8, we retained only one gene from the pair.

#### Preprocessing and filtering of SCNA data

We split each chromosome into contiguous segments, each spanning 1 Mb. For each segment, the mean signal intensity was calculated, providing an average measure of copy number changes with positive values indicating an amplification and a negative value indicating a deletion. We retained the segments with the highest 70% variance, focusing our analysis on regions with the most substantial variation in copy number. Certain tumors exhibit extensive copy number alterations, where alterations might extend across an entire chromosomal arm or a large segment of a chromosome. As a result, consecutive segments within these regions often show correlated intensities. We performed filtering in two steps: (1) for regions where SCNA value was consistent across all samples, we kept only the first segment, and (2) we performed correlation-based pruning so that no segments had pairwise correlation *R*^2^ ≥ 0.8. To make analysis robust, the measure of SCNA was quantile normalized to approximately follow a normal distribution.

#### Somatic mutations in driver genes of LUAD

We downloaded the somatic mutation data from https://www.cbioportal.org/datasets while exome sequencing of thirteen driver genes for LUAD, including *ALK, EGFR, KEAP1, KRAS, NOTCH1, NTRK1, PIK3CA, RB1, RET, ROS1, SMARCA4, STK11*, and *TP53*. For each driver gene, a subject was marked as mutated if a potentially function mutation was detected in the following categories: missense mutation, nonsense mutation, nonstop mutation, frame shift insertion, in frame insertion, and frame shift deletion.

#### Adjusting for purity in analysis

Purity of a tumor measures the fraction of tumor cells in the tumor sample and is a crucial confounding factor when analyzing the relation between SCNA and expression, as was empirically demonstrated. We adjusted for purity in our analysis by using purity as a covariate. We downloaded three purity metrics for each subject from TCGA^62^, inferred based on SCNAs, SNVs, and DNA methylation, and used the median in our analysis.

### Gut Metabolites and Metagenomic Data in Japanese Colorectal Cancer Cohort

Meta-data, gut metabolite data, and genus-level taxonomic abundances were obtained from Yachida et al. (2019)^22^. Our analysis included 347 subjects (127 healthy controls, 67 MP/stage 0 CRC patients, 69 stage 1/II CRC patients, and 54 stage III/IV CRC patients, and 30 patients with history of colorectal surgery) with both gut metabolite and metagenomic data. To ensure robustness, we applied quantile normalization to each metabolite. We selected 225 genera for analysis that were present in more than 70% of samples, ensuring sufficient variability for statistical power.

### Simulations

Given *M* biological traits, let *Θ* = (*θ*_*ij*_) ∈ ℝ^*M*×*M*^ represent the precision matrix of the baseline network. In our simulations, the baseline network was structured with 10 hubs, each connected to four nodes. The diagonal elements of *Θ* were set to 10, while *θ*_*ij*_ = 2.5 if an edge existed between nodes (*i, j*), and 0 otherwise. Under this configuration, the baseline or the population average PCC was *ρ*_*ij*_ = 0.25.

We chose to model common SNPs as genomic factors, with minor allele frequencies (MAF) uniformly distributed in the range [0.05, 0.5]. For *N* samples, we simulated *T* correlated genetic variants by grouping them into 5 non-overlapping blocks and we utilized a Toeplitz structure to model the correlation among the variants in each block with correlation coefficient *ρ* = 0.5. Let *g*_*nt*_ represent the number of the minor alleles for variant *t* and sample *n*. In our simulations, we allowed 10 edges, each originating from one hub (**Figure 2a**), to be modulated by one individual SNP. For the pair with a signal, we assumed a trend effect model:

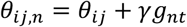

to characterize the relationship between the subject-specific partial covariance *θ*_*ij,n*_ (and thus PCC *ρ*_*ij,n*_) and host genotype. We considered three effect sizes *γ* = 0.25, 0.30, and 0.35, which correspond to different PCCs across three genotypic groups. For example, when *γ* = 0.25, PCC = 0.25, 0.50, and 0.75 for genotypes 0, 1, and 2, respectively; when *γ* = 0.35, PCC = 0.25, 0.60, and 0.95 (Figure 2B), respectively. Thus, a large effect size *π* corresponds a large difference of PCCs in three genotypic groups. Additionally, selecting these effect sizes ensured that the subject-specific PCC remained within [-1,1] and that the precision matrix remained positive definite. Finally, we generated *M* biological traits for each *n*^*th*^ sample from a multivariate normal distribution with a covariance matrix: 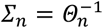.

For each configuration of simulation parameters and a tuning parameter λ, we conducted 50 repeats to assess whether the estimated FDR aligned with the true FDR, and to evaluate sensitivity, defined as the proportion of correctly detected signals. Since the false and correct detections were precisely known, the true FDR could be directly calculated.

## Acknowledgement

The project was supported by the NIH Intramural Research Program. This study utilized the high-performance computational capabilities of the Biowulf Linux cluster at the National Institutes of Health, Bethesda, MD. (http://biowulf.nih.gov).

## Author Contributions

S.A., P.S.A, and J.S conceived the project. S.A., S.L, X.W., X.H., P.S.A, and J.S contributed to algorithm development and data analysis. S.A., P.S.A, and J.S. draft the manuscript. S.A., G.H.O., E.V., M.T.L., W.Z., X.R.Y., M.L.R, K.B, B.Z., K.Y., S.C.M, C.A., T.Z., P.S.A., J.S. interpreted results. All authors reviewed the manuscript.

## Competing interests

The authors declare no conflict of interest.

## Data availability

For PLCO, the oral microbiome sequencing data are available at the Sequence Read Archive (SRA) under project number PRJNA801882 with limited metadata (https://www.ncbi.nlm.nih.gov/sra/). The meta data, gene expression, SCNA, and somatic mutation data are downloaded from TCGA at https://www.cbioportal.org/datasets. For the CRC in the Japanese cohort, the genera abundance data and the gut metabolite data are extracted from Supplementary Table S7-2 and Supplementary Table S13, respectively of Yachida et al. (2019).

## Code availability

The developed software and the test data are available from GitHub at https://github.com/samuelanyaso/GFBioNet.

## Notes

### Competing Interest Statement

The authors have declared no competing interest.

## References

1. Lauritzen, S. Graphical Models, (Oxford: Oxford University Press, 1996).

2. Yuan, M. & Lin, Y. Model selection and estimation in the Gaussian graphical model. Biometrika 94, 19–35 (2007).

3. Friedman, J., Hastie, T. & Tibshirani, R. Sparse inverse covariance estimation with the graphical lasso. Biostatistics 9, 432–441 (2008).

4. Zhao, H. & Duan, Z.H. Cancer Genetic Network Inference Using Gaussian Graphical Models. Bioinform Biol Insights 13, 1177932219839402 (2019).

5. Jiang, D. et al. Microbiome Multi-Omics Network Analysis: Statistical Considerations, Limitations, and Opportunities. Front Genet 10, 995 (2019).

6. Faust, K. et al. Microbial co-occurrence relationships in the human microbiome. PLoS Comput Biol 8, e1002606 (2012).

7. Xu, Y., Jiang, H. & Jiang, W. Extended graphical lasso for multiple interaction networks for high dimensional omics data. PLoS Comput Biol 17, e1008794 (2021).

8. Krumsiek, J., Suhre, K., Illig, T., Adamski, J. & Theis, F.J. Gaussian graphical modeling reconstructs pathway reactions from high-throughput metabolomics data. BMC Syst Biol 5, 21 (2011).

9. Feng, L. et al. Proteome-wide data analysis reveals tissue-specific network associated with SARS-CoV-2 infection. J Mol Cell Biol 12, 946–957 (2020).

10. Mazumder, R. & Hastie, T. The graphical lasso: New insights and alternatives. Electronic Journal of Statistics 6, 2125–2149 (2012).

11. Meinshausen, N. & Bühlmann, P. High-dimensional graphs and variable selection with the Lasso. Annals of Statistics 34, 1436–1462 (2006).

12. Magwene, P.M. & Kim, J. Estimating genomic coexpression networks using first-order conditional independence. Genome Biol 5, R100 (2004).

13. Wille, A. & Buhlmann, P. Low-order conditional independence graphs for inferring genetic networks. Stat Appl Genet Mol Biol 5, Article1 (2006).

14. Faming Liang, Q.S., Peihua Qiu. An Equivalent Measure of Partial Correlation Coefficients for High-Dimensional Gaussian Graphical Models. Journal of the American Statistical Association 110, 1248–1265 (2014).

15. Danaher, P., Wang, P. & Witten, D.M. The joint graphical lasso for inverse covariance estimation across multiple classes. J R Stat Soc Series B Stat Methodol 76, 373–397 (2014).

16. Shojaie, A. Differential Network Analysis: A Statistical Perspective. Wiley Interdiscip Rev Comput Stat 13(2021).

17. Tan, K.M., London, P., Mohan, K., Lee, S.I., Fazel, M. & Witten, D. Learning Graphical Models With Hubs. J Mach Learn Res 15, 3297–3331 (2014).

18. Tan, K.M., Witten, D. & Shojaie, A. The cluster graphical lasso for improved estimation of Gaussian graphical models. Comput Stat Data Anal 85, 23–36 (2015).

19. Deng, W., Zhang, K., Liu, S., Zhao, P.X., Xu, S. & Wei, H. JRmGRN: joint reconstruction of multiple gene regulatory networks with common hub genes using data from multiple tissues or conditions. Bioinformatics 34, 3470–3478 (2018).

20. Yazar, S. et al. Single-cell eQTL mapping identifies cell type-specific genetic control of autoimmune disease. Science 376, eabf3041 (2022).

21. Zhang, J. & Li, Y. High-Dimensional Gaussian Graphical Regression Models with Covariates. J Am Stat Assoc 118, 2088–2100 (2023).

22. Yachida, S. et al. Metagenomic and metabolomic analyses reveal distinct stage-specific phenotypes of the gut microbiota in colorectal cancer. Nat Med 25, 968–976 (2019).

23. Vogtmann, E. et al. The Oral Microbiome and Lung Cancer Risk: An Analysis of 3 Prospective Cohort Studies. J Natl Cancer Inst 114, 1501–1510 (2022).

24. Cancer Genome Atlas Research, N. Comprehensive molecular profiling of lung adenocarcinoma. Nature 511, 543–50 (2014).

25. Rina Foygel, M.D. Extended Bayesian Information Criteria for Gaussian Graphical Models. in NIPS (2010).

26. Van de Geer, S., Bühlmann, P., Ritov, Y. & Dezeure, R. On Asymptotically Optimal Confidence Regions and Tests for High-Dimensional Models. Annals of Statistics 42, 1166–1202 (2014).

27. Javanmard, A. & Montanari, A. Confidence Intervals and Hypothesis Testing for High-Dimensional Regression. Journal of Machine Learning Research 15, 2869–2909 (2014).

28. Breheny, P.J. Marginal false discovery rates for penalized regression models. Biostatistics 20, 299–314 (2019).

29. Henao, J.D. et al. Multi-omics regulatory network inference in the presence of missing data. Brief Bioinform 24 (2023).

30. Louis, P., Hold, G.L. & Flint, H.J. The gut microbiota, bacterial metabolites and colorectal cancer. Nat Rev Microbiol 12, 661–72 (2014).

31. Moskovitz, J., Yim, M.B. & Chock, P.B. Free radicals and disease. Arch Biochem Biophys 397, 354–9 (2002).

32. Wallace, H.M., Fraser, A.V. & Hughes, A. A perspective of polyamine metabolism. Biochem J 376, 1–14 (2003).

33. Li, C. et al. Gut microbiome and metabolome profiling in Framingham heart study reveals cholesterol-metabolizing bacteria. Cell 187, 1834–1852 e19 (2024).

34. Yu, J. et al. Metagenomic analysis of faecal microbiome as a tool towards targeted non-invasive biomarkers for colorectal cancer. Gut 66, 70–78 (2017).

35. Goodrich, J.K. et al. Human genetics shape the gut microbiome. Cell 159, 789–99 (2014).

36. Kurilshikov, A. et al. Large-scale association analyses identify host factors influencing human gut microbiome composition. Nat Genet 53, 156–165 (2021).

37. Kurtz, Z.D., Muller, C.L., Miraldi, E.R., Littman, D.R., Blaser, M.J. & Bonneau, R.A. Sparse and compositionally robust inference of microbial ecological networks. PLoS Comput Biol 11, e1004226 (2015).

38. Davies, C.S. et al. Immunogenetic variation shapes the gut microbiome in a natural vertebrate population. Microbiome 10, 41 (2022).

39. Blekhman, R. et al. Host genetic variation impacts microbiome composition across human body sites. Genome Biol 16, 191 (2015).

40. Kubinak, J.L. et al. MHC variation sculpts individualized microbial communities that control susceptibility to enteric infection. Nat Commun 6, 8642 (2015).

41. Tett, A., Pasolli, E., Masetti, G., Ercolini, D. & Segata, N. Prevotella diversity, niches and interactions with the human host. Nat Rev Microbiol 19, 585–599 (2021).

42. Simon-Soro, A. & Mira, A. Solving the etiology of dental caries. Trends Microbiol 23, 76–82 (2015).

43. Baty, J.J., Stoner, S.N. & Scoffield, J.A. Oral Commensal Streptococci: Gatekeepers of the Oral Cavity. J Bacteriol 204, e0025722 (2022).

44. Ricklin, D., Hajishengallis, G., Yang, K. & Lambris, J.D. Complement: a key system for immune surveillance and homeostasis. Nat Immunol 11, 785–97 (2010).

45. Cargill, J.S., Scott, K.S., Gascoyne-Binzi, D. & Sandoe, J.A.T. Granulicatella infection: diagnosis and management. J Med Microbiol 61, 755–761 (2012).

46. Erwin, A.L. & Smith, A.L. Nontypeable Haemophilus influenzae: understanding virulence and commensal behavior. Trends Microbiol 15, 355–62 (2007).

47. Scher, J.U. et al. Expansion of intestinal Prevotella copri correlates with enhanced susceptibility to arthritis. Elife 2, e01202 (2013).

48. Hwang, J.H., Yoon, J., Cho, Y.H., Cha, P.H., Park, J.C. & Choi, K.Y. A mutant KRAS-induced factor REG4 promotes cancer stem cell properties via Wnt/beta-catenin signaling. Int J Cancer 146, 2877–2890 (2020).

49. Sun, S. et al. REG4 is an indicator for KRAS mutant lung adenocarcinoma with TTF-1 low expression. J Cancer Res Clin Oncol 145, 2273–2283 (2019).

50. Zheng, H.C., Xue, H. & Zhang, C.Y. REG4 promotes the proliferation and anti-apoptosis of cancer. Front Cell Dev Biol 10, 1012193 (2022).

51. Adinolfi, S. et al. The KEAP1-NRF2 pathway: Targets for therapy and role in cancer. Redox Biol 63, 102726 (2023).

52. Taguchi, K., Motohashi, H. & Yamamoto, M. Molecular mechanisms of the Keap1-Nrf2 pathway in stress response and cancer evolution. Genes Cells 16, 123–40 (2011).

53. Shibata, T. et al. Cancer related mutations in NRF2 impair its recognition by Keap1-Cul3 E3 ligase and promote malignancy. Proc Natl Acad Sci U S A 105, 13568–73 (2008).

54. Stegle, O., Parts, L., Piipari, M., Winn, J. & Durbin, R. Using probabilistic estimation of expression residuals (PEER) to obtain increased power and interpretability of gene expression analyses. Nat Protoc 7, 500–7 (2012).

55. Sun, W. et al. The association between copy number aberration, DNA methylation and gene expression in tumor samples. Nucleic Acids Res 46, 3009–3018 (2018).

56. Albert, F.W., Bloom, J.S., Siegel, J., Day, L. & Kruglyak, L. Genetics of trans-regulatory variation in gene expression. Elife 7(2018).

57. Joo, J.W., Sul, J.H., Han, B., Ye, C. & Eskin, E. Effectively identifying regulatory hotspots while capturing expression heterogeneity in gene expression studies. Genome Biol 15, r61 (2014).

58. Tibshirani, R. Regression shrinkage and selection via the Lasso. Journal of the Royal Statistical Society Series B-Methodological 58, 267–288 (1996).

59. Zou, H. & Hastie, T. Regularization and variable selection via the elastic net. Journal of the Royal Statistical Society Series B-Statistical Methodology 67, 301–320 (2005).

60. Liu, H., Roeder, K. & Wasserman, L. Stability Approach to Regularization Selection (StARS) for High Dimensional Graphical Models. Adv Neural Inf Process Syst 24, 1432–1440 (2010).

61. Vogtmann E, H.X., Yu G, Purandare V, Hullings AG, Shao D, Wan Y, Li S, Dagnall CL, Jones K, Hicks BD, Hutchinson A, Caporaso JG, Wheeler W, Sandler DP, Beane Freeman LE, Liao LM, Huang WY, Freedman ND, Caporaso N, Sinha R, Gail MH, Shi J, Abnet CC. . The human oral microbiome and risk of lung cancer: An analysis of three prospective cohort studies. . JAMA Oncology 2022;ONC21-3241. (2022).

62. Aran, D., Sirota, M. & Butte, A.J. Systematic pan-cancer analysis of tumour purity. Nat Commun 6, 8971 (2015).

